# Identification of fungal limonene-3-hydroxylases for biotechnological menthol production

**DOI:** 10.1101/2020.11.25.399378

**Authors:** Florence M. Schempp, Ingmar Strobel, Maria M. W. Etschmann, Elena Bierwirth, Johannes Panten, Hendrik Schewe, Jens Schrader, Markus Buchhaupt

## Abstract

More than 30,000 tons of menthol are produced every year as a flavor and fragrance compound or as medical component. So far, only extraction from plant material or chemical synthesis is possible. A sustainable alternative approach for menthol production could be a biotechnological-chemical two-step conversion, starting from (+)-limonene, which is a side product of the citrus processing industry. The first step requires a limonene-3-hydroxylase (L3H) activity that specifically catalyzes hydroxylation of limonene at carbon atom 3. Several protein engineering strategies already attempted to create limonene-3-hydroxylases from bacterial cytochrome P450 monooxygenases (CYPs or P450s), which can be efficiently expressed in bacterial hosts. However, their regiospecificity is rather low, if compared to the highly selective L3H enzymes from the biosynthetic pathway towards menthol in *Mentha* species. The only naturally occurring limonene-3-hydroxylase activity identified in microorganisms so far, was reported for a strain of the black yeast-like fungus *Hormonema* sp. in South Africa.

We have discovered further fungi that can catalyze the intended reaction and identified potential CYP-encoding genes within the genome sequence of one of the strains. Using heterologous gene expression and biotransformation experiments in yeasts, we were able to identify limonene-3-hydroxylases from *Aureobasidium pullulans* and *Hormonema carpetanum*. Further characterization of the *A. pullulans* enzyme demonstrated its high stereospecificity and regioselectivity, its potential for limonene-based menthol production and its additional ability to convert α-and β-pinene to verbenol and pinocarveol, respectively.

**Importance:** (−)-Menthol is an important flavor and fragrance compound and furthermore has medicinal uses. To realize a two-step synthesis starting from renewable (+)-limonene, a regioselective limonene-3-hydroxylase enzyme is necessary. We identified enzymes from two different fungi, which catalyze this hydroxylation reaction and represent an important module for the development of a biotechnological process for (−)-menthol production from renewable (+)-limonene.

## Introduction

(−)-Menthol is a monocyclic monoterpene alcohol with an annual production volume of more than 30,000 tons (1). It is used as a flavor and fragrance compound, as cosmetics additive and as ingredient of pharmaceutical products, e.g. for the treatment of colds or burns (2). Menthol can either be extracted from the essential oil of *Mentha* plants or produced by various chemical processes. The technical synthesis of (−)-menthol is currently carried out using different processes at Symrise/Lanxess (from *m*-cresol), Takasago (from myrcene) and BASF (from citral) (2, 3). However, no biotechnological process for (−)-menthol production has been established so far. Since the product yields of *Mentha* plantations are strongly dependent on changing climatic factors and chemical synthesis is partly based on limited fossil resources, biotechnological production processes based on renewable raw materials are an attractive alternative. A sustainable menthol synthesis route could be a biotechnological-chemical two-step conversion (4), starting from the renewable raw material limonene. The conversion of limonene into the intermediate requires specific hydroxylase enzymes. In the case of (−)-limonene a limonene-5-hydroxylase (L5H) would be necessary to form *trans*-*p*-1,8-menthadien-5-ol, whereas (+)-limonene has to be hydroxylated at carbon atom 3 by a limonene-3-hydroxylase (L3H) to produce *trans*-isopiperitenol (Figure 1). *trans*-Isopiperitenol, and presumably also *trans*-*p*-1,8-menthadien-5-ol, can then be further transformed into (−)-menthol by chemical hydrogenation (4, 5). Many protein engineering strategies have already attempted to create limonene-3-hydroxylases with high regioselectivity and stereospecificity from bacterial cytochrome P450 monooxygenase (CYPs or P450s) (6–8). However, while these enzymes can be efficiently expressed in bacterial hosts, their product selectivity is limited (75 % *trans*-isopiperitenol out of (+)-limonene at best with formation of several side products) (7).

**Figure 1:**
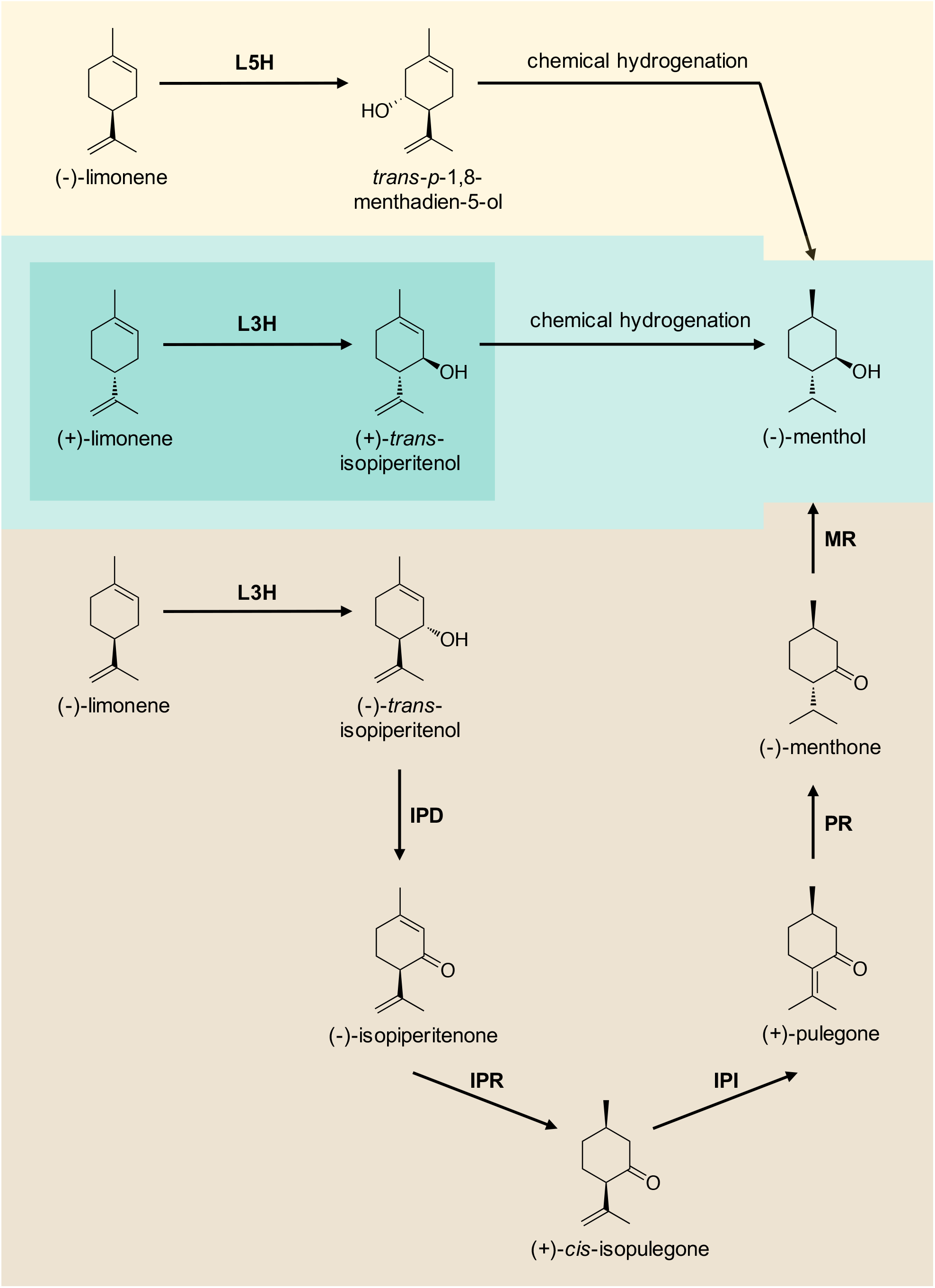
Biotechnological-chemical processes for (−)-menthol production (upper part – yellow and green) vs. native pathway in peppermint plant (*Mentha* x*piperita*) (lower part – brown, adapted from (9)). L5H: limonene-5-hydroxylase, L3H: limonene-3-hydroxylase, IPD: (−)-*trans*-isopiperitenol dehydrogenase, IPR: (−)-isopiperitenone reductase, IPI: (+)-*cis*-isopulegone isomerase, PR: (+)-pulegone reductase, MR: (−)-menthone reductase.

In the peppermint plant *Mentha x piperita,* (−)-menthol is produced from (−)-limonene via a six-step metabolic pathway (9) (Figure 1). The first enzyme of this pathway is a selective limonene-3-hydroxylase which is present in two slightly different forms encoded by CYP71D13 (PM17) and CYP71D15 (PM2) (10). As with other enzymes of the menthol pathway (2), several studies have been conducted to express PM17 heterologously in microbial hosts (11, 12), but resulted in rather low productivities. Besides their occurrence in *Mentha* plants (2, 10, 13), the only other limonene hydroxylase activity with clear selectivity for carbon atom 3 was described for a strain of the black yeast-like fungus *Hormonema* sp. (UOFS Y-0067) (4). The genus *Hormonema* belongs to the class of Dothideomycetes, which in turn forms a subgroup of Ascomycota. Many Dothideomycetes live as endophytes or epiphytes in or on plants, or degrade dead plant material as saprobionts.(14) The cells of *Hormonema* sp. UOFS Y-0067 were also found to hydroxylate α- and β-pinene. In the respective report, van Dyk and colleagues (1998) already suggested the potential use of the fungus for (−)-menthol production from (+)-limonene, but the results were not reproducible and the obtained product concentrations varied greatly. In addition, the responsible enzyme remained unknown. Therefore, they considered it unlikely “that it will ever be possible to base a commercial biotransformation process on an organism like this *Hormonema* sp.”(4)

In order to identify stereo- and regioselective limonene-3-hydroxylases in *Hormonema* or other fungi, we tested different related fungi for the respective conversion. Candidate CYP enzymes were identified in the fungal genomes and by gene comparison and heterologous expression of twenty different CYP sequences in yeasts, we identified a limonene-3-hydroxylase from *Aureobasidium pullulans* and also from *Hormonema carpetanum*. Subsequently, the enzyme of *A. pullulans* was further characterized in detail regarding substrate and product selectivity.

## Results

### Hydroxylation capability of different Dothideomycetes

In order to investigate if the limonene-3-hydroxylase activity of *Hormonema* sp. UOFS Y-0067 (4) is a common ability in the class of Dothideomycetes, several fungi, including *Hormonema carpetanum*, *Aureobasidium pullulans*, *Neofusicoccum parvum*, *Ochroconis constricta* and *Parastagonospora nodorum,* were tested for (+)-limonene hydroxylation. *Hormonema carpetanum* and *Aureobasidium pullulans* were found to convert limonene into *trans*-isopiperitenol, while *A. pullulans* showed the highest *trans*-isopiperitenol production over time (Figure 2 A). Further analyses confirmed that *A. pullulans* produces *trans*-isopiperitenol as main product in a limonene biotransformation reaction (Figure 2 B - D). Therefore, this strain was used to search for the respective hydroxylase enzyme.

**Figure 2:**
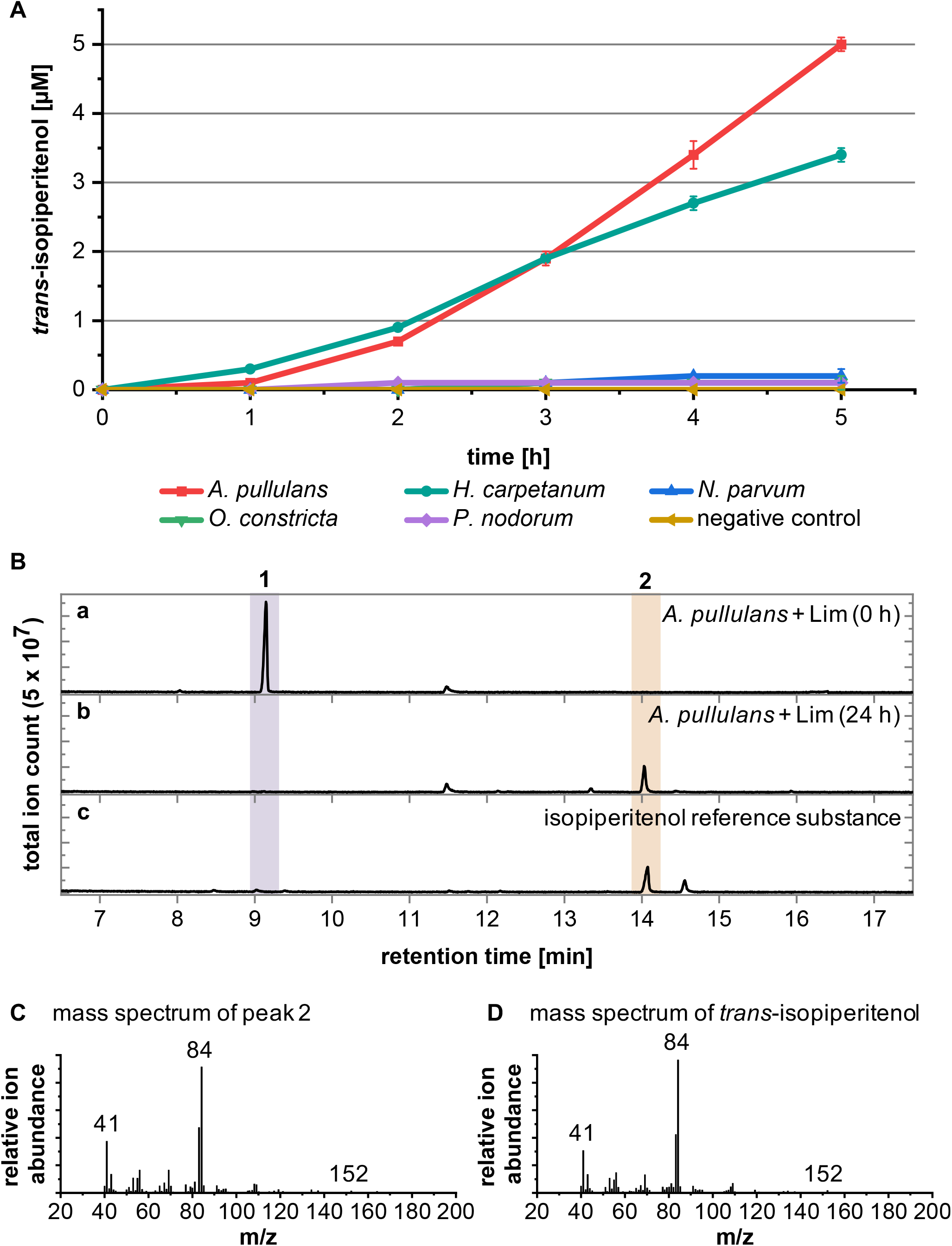
*trans*-Isopiperitenol production with different Dothideomycetes. (A) *trans*-Isopiperitenol production over time of different Dothideomycetes. Culture samples of different fungi were taken over a period of 5 hours (once per hour) after addition of 1 g L^−1^(+)-limonene. Samples were extracted with ethyl acetate and the *trans*-isopiperitenol amount was determined by GC-MS analysis. Negative control: medium without cells. The data points and error bars represent the mean values and standard deviations of three biological replicates (n = 3). (B) GC-MS analysis of biotransformation of (+)-limonene by *A. pullulans*. Chromatogram of ethyl acetate extract from (a) *A. pullulans* culture after addition of 1 g L^−1^ limonene (t = 0 h) is compared to (b) *A. pullulans* culture 24 h after addition of limonene, and (c) isopiperitenol reference substance (2:1-mixture of (+)-*trans-/cis*-isopiperitenol). *trans*-Isopiperitenol (peak 2, retention time (Rt.) 14 min) was identified as the main product of (+)-limonene (peak 1) biotransformation by comparison of retention time and (C) mass-spectrum to those of (D) the isopiperitenol reference substance.

### Identification of fungal limonene-3-hydroxylase enzymes

Since preliminary experiments with specific enzyme inhibitors had shown that the responsible protein is most probably a member of the cytochrome P450 monooxygenase family (data not shown), we focused on such enzymes. According to the NCBI database, the genome sequence of *A. pullulans* CBS 100280 (EXF-150) (15) contains 51 genes encoding putative CYP proteins. These annotated CYPs from *A. pullulans* were assigned into different families based on their protein sequence and named accordingly by David Nelson (16) (Figure 3). The individual proteins were clustered by means of multiple sequence alignment (Figure 3). Sixteen of the 51 putative *A. pullulans* CYPs were chosen for expression in *Saccharomyces cerevisiae,* so that candidates from the different subclusters of the phylogenetic tree were included (Figure 3 – blue dots, Table S1).

**Figure 3:**
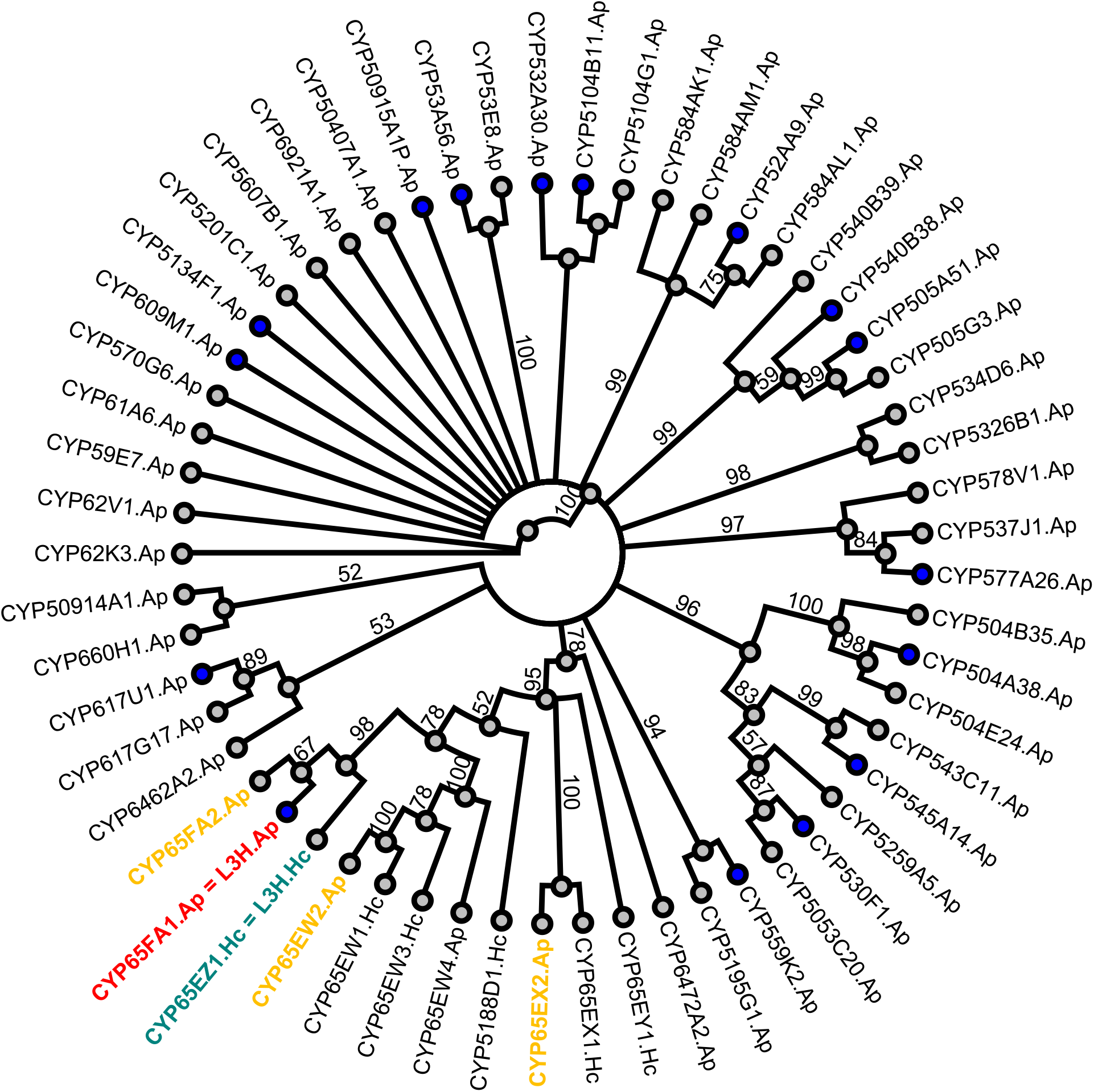
Clustering of cytochrome P450 monooxygenases of *A. pullulans* CBS 100280 (EXF-150) and of *H. carpetanum* CBS 115712. (only selected ones with the highest identity to L3H.Ap). The proteins annotated as cytochrome P450 monooxygenases were analyzed for their amino acid sequence identity and subdivided into protein families. CYP candidate genes of *A. pullulans* chosen for expression in *S. cerevisiae* are marked with blue dots. Limonene-3-hydroxylase enzymes from *A. pullulans* (L3H.Ap = CYP65FA1) and *H. carpetanum* (L3H.Hc = CYP65EZ1) as well as other tested CYP65 candidates from *A. pullulans* are marked in red, green or yellow, respectively. For accession numbers see Table S1. Clustering was performed with the software *Geneious* using default settings (ClustalW alignment, Geneious tree builder, resampling method: bootstrap, number of replicates: 1,000). Branch labels: Consensus support [%].

In addition to the potential CYP-encoding sequences, a gene encoding a putative CYP reductase (CPR) enzyme was identified in the genome of *A. pullulans* (Table S1), by comparing protein sequences of known CYP reductases from other fungi with the genome of *A. pullulans*.

In an initial investigation, *S. cerevisiae* strains harboring the different CYP genes together with the putative CPR were tested in a (+)-limonene biotransformation experiment. One strain led to a clearly increased *trans*-isopiperitenol concentration compared to the background product level of the strain containing the empty vectors (Figure 4, Figure S1). Therefore, we proposed the respective protein CYP65FA1 of *A. pullulans* (Ap) to be a limonene-3-hydroxylase, hereafter referred to as CYP65FA1.Ap or L3H.Ap. In addition, it was found that also without the co-expression of the putative CYP reductase, *trans*-isopiperitenol was produced with the respective *S. cerevisiae* strain, indicating that L3H.Ap can also work together with a native yeast CPR.

**Figure 4:**
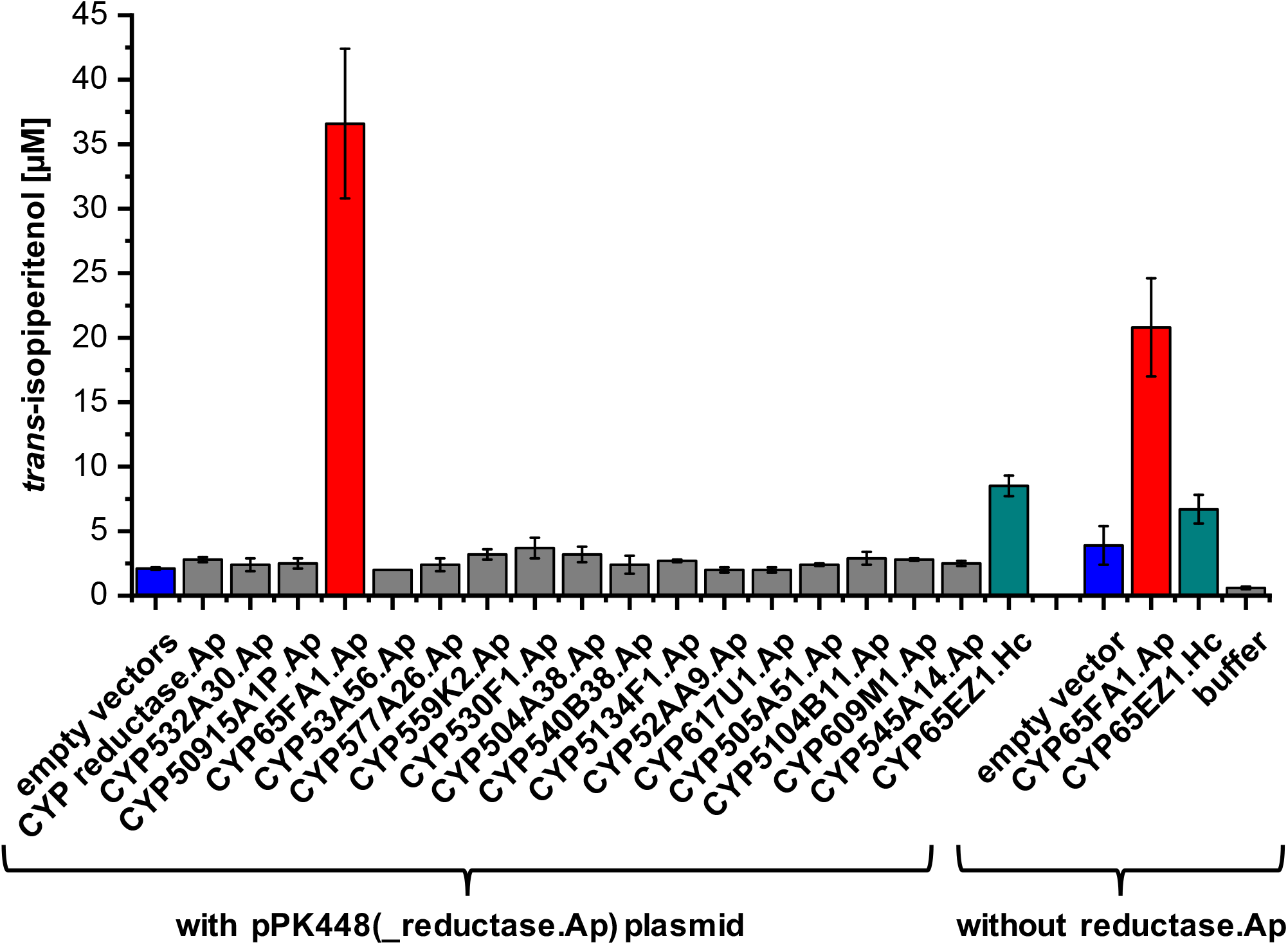
*trans*-Isopiperitenol concentrations achieved in (+)-limonene biotransformation experiment with different *S. cerevisiae* CEN-PK2-1C strains. Concentrated cell suspensions were used. empty vector(s): yeast strain harboring empty pPK245 (and pPK448) vectors, CYP reductase.Ap: cells expressing putative CYP reductase gene from *A. pullulans* CBS 100280 (EXF-150) from pPK448 vector and containing empty pPK245, CYP532A30.Ap – CYP65EZ1.Hc: cells expressing different putative CYPs from *A. pullulans* CBS 100280 (EXF-150, .Ap) or *H. carpetanum* CBS 115712 (.Hc) from pPK245 vector together with putative CYP reductase from *A. pullulans* from pPK448 plasmid (left side) or without reductase.Ap (right side). *trans*-Isopiperitenol concentrations were calculated from peak areas, which were determined by GC-MS analysis, and normalized to the peak area of the internal standard (3-carene) and the number of cells corresponding to 150 OD units. The data points and error bars represent the mean values and standard deviations of three biological replicates (n = 3).

In order to identify further limonene-3-hydroxylase enzymes in the genome of *A. pullulans* and *H. carpetanum*, the second fungus for which L3H activity was detected in the previous experiments, additional putative CYP genes of both fungi were selected based on amino acid sequence identity to CYP65FA1.Ap. Thereby, CYP65FA2, CYP65EX2 and CYP65EW2 from *A. pullulans* (Ap) and CYP65EZ1 from *H. carpetanum* (Hc) were chosen for further tests (Figure 3 – yellow and green font). The CYP65EZ1.Hc gene sequence was identified by analysis of partial genome data of *H. carpetanum* CBS 115712. The amino acid sequence identity between CYP65FA1.Ap (L3H.Ap) and CYP65FA2.Ap, CYP65EX2.Ap, CYP65EW2.Ap or CYP65EZ1.Hc, is about 53 %, 33 %, 41 % and 52 %, respectively. Whereas no limonene conversion could be observed in the biotransformation experiments with CYP65FA2.Ap, CYP65EX2.Ap and CYP65EW2.Ap (data not shown), *trans*-isopiperitenol could be detected in biotransformation samples with the yeast strain expressing CYP65EZ1 from *H. carpetanum* (Figure 4), further referred to as L3H.Hc. However, the concentrations achieved were less than a third of the value of the strains harboring the L3H.Ap gene. Therefore, the L3H enzyme of *A. pullulans* was chosen for further characterization.

### Further characterization of limonene-3-hydroxylase abilities of L3H.Ap

In order to obtain higher protein amounts to further characterize the limonene-3-hydroxylase enzyme CYP65FA1 of *A. pullulans* (L3H.Ap), we tried to express the protein and several variants in *E. coli*. After all attempts to express the protein in a functional form with *E. coli* failed, we tested expression in the eukaryotic host *Pichia pastoris* (17). For this purpose, two different vector systems were applied. Whereas one of the used vectors is an episomal plasmid with constitutive expression (pBSYA1Z), the pPICZA vector is integrated into the genome of *P. pastoris* and expression of the CYP is induced by methanol addition. As negative control, the cells were transformed with the respective empty expression vector. Furthermore, PM17 from *Mentha x piperita,* codon-optimized for *P. pastoris* (12), was used as a reference. Since it has been observed that the L3H.Ap enzyme is functional in *S. cerevisiae* with the native yeast CPR, the putative CYP reductase of *A. pullulans* was not co-expressed in *P. pastoris.* The experiments showed that with both vector systems higher amounts of *trans*-isopiperitenol were produced from (+)-limonene with L3H.Ap of *A. pullulans* than with PM17 from *Mentha* x *piperita* (Figure 5 A, Figure S2). The maximum *trans*-isopiperitenol concentration obtained was around 300 μM (45 mg L^−1^) in the concentrated yeast cell suspension with constitutive gene expression from the episomal vector.

**Figure 5:**
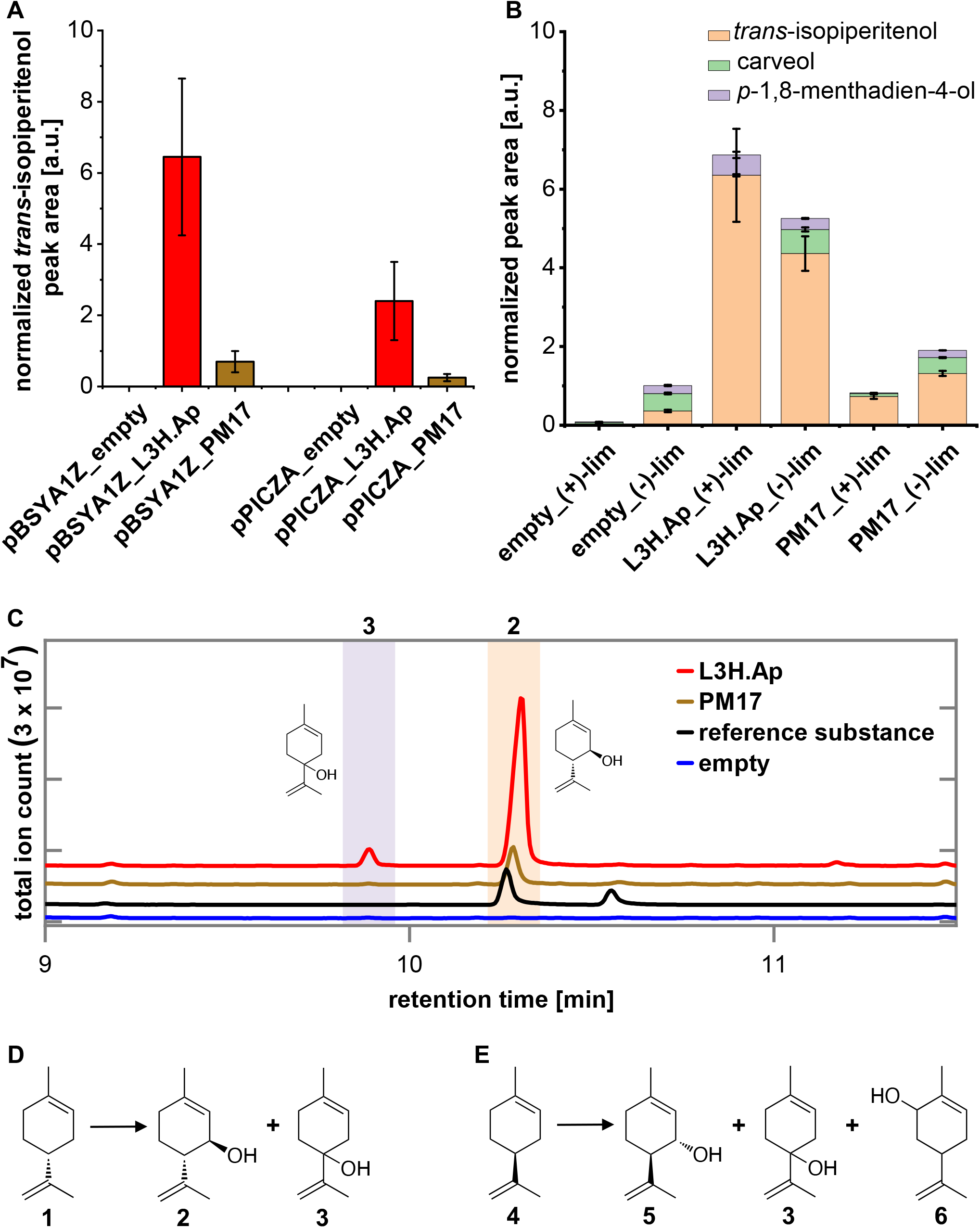
Hydroxylation of limonene with *P. pastoris* X-33 strains expressing L3H enzymes. (A) *trans*-Isopiperitenol peak areas achieved in (+)-limonene biotransformation experiment with different *P. pastoris* X-33 strains (concentrated cell suspensions). pBSYA1Z: episomal, constitutive gene expression, pPICZA: genome integrated, methanol-induced gene expression. empty: *P. pastoris* with empty expression vector, L3H.Ap: with CYP65FA1 of *A. pullulans*, PM17: with limonene-3-hydroxylase enzyme of *Mentha x piperita.* (B) Product peak areas achieved in biotransformation experiment with (+)-or (−)-limonene with *P. pastoris* X-33 strains (concentrated cell suspensions) harboring empty pBSYA1Z plasmid or expressing limonene-3-hydroxylase enzymes from *A. pullulans* (L3H.Ap) or *Mentha* x *piperita* (PM17) from pBSYA1Z backbone. While with (−)-limonene several different products were formed with all strains (see Figure S3), only peak areas of the main (by-)products are shown for comparison. Chromatograms see Figure 5 C, Figure S2 and Figure S3. In (A) and (B) peak areas were determined by GC-MS analysis, and normalized to the peak are of the internal standard (3-carene) and the number of cells corresponding to (A) 150 or (B) 225 OD units. The data points and error bars represent the mean values and standard deviations of six (A: n = 6) or three (B: n = 3) biological replicates. (C) GC-MS analysis of (+)-limonene biotransformation experiment shown in Figure 5 B. Chromatogram of the negative control with the empty pBSYA1Z expression plasmid (blue) is compared to a strain harboring PM17 from *Mentha* x *piperita* (brown) or L3H.Ap (red), and isopiperitenol reference substance (2:1-mixture of (+)-*trans-/cis*-isopiperitenol) (black). *trans*-Isopiperitenol (peak 2, Rt. 10.3 min) was identified as the main product (> 90 %) of (+)-limonene biotransformation. As side product (< 10 %) *p-*1,8-menthadien-4-ol (peak 3, Rt. 9.8 min) was formed by cells expressing L3H.Ap. (D, E) Substrates and main products of bioconversion reactions catalyzed by L3H.Ap. 1: (+)-limonene, 2: (+)-*trans*-isopiperitenol, 3: *p-*1,8-menthadien-4-ol, 4: (−)-limonene, 5: (−)-*trans*-isopiperitenol, 6: carveol (MS spectrum see Figure S4).

Apart from *trans*-isopiperitenol, *p*-1,8-menthadien-4-ol could be detected in the limonene biotransformation reactions with L3H.Ap (Figure 5 B - D, Figure S2). However, the peak area of produced *trans*-isopiperitenol is around 10 times larger than the *p*-1,8-menthadien-4-ol peak area. Small *p*-1,8-menthadien-4-ol peaks could also be identified in the limonene biotransformation experiments with *A. pullulans* cultures (Figure 2, retention time (Rt.) 13.4 min) and *S. cerevisiae* cells harboring the fungal L3H enzyme (Figure S1, Rt. 9.8 min). Further oxidation of the formed *trans*-isopiperitenol or *p*-1,8-menthadien-4-ol was not observed (Figure S2).

When comparing both limonene enantiomers, higher product peaks were obtained with (+)-limonene than with (−)-limonene in a biotransformation experiment with the L3H.Ap-expressing strain (Figure 5 B, Figure S2, Figure S3). PM17 showed an inverse preference (Figure 5 B, Figure S2, Figure S3).

Unexpectedly, when (−)-limonene was used as substrate several different products were formed (Figure S3). However, this could be observed with all strains, harboring the empty vector, L3H.Ap or PM17 from *Mentha* x *piperita,* to a similar extent. Only the production of *trans*-isopiperitenol was clearly increased compared to the negative control when the fungal or plant L3H was present (Figure 5 B).

Alteration of different parameters to optimize the biotransformation process yielded higher *trans*-isopiperitenol amounts and in relation to this reduced side product formation. Beneficial modifications were the alteration of the pH of the biotransformation buffer from 6.0 to 7.4 or 8.0 (Figure 6 A), and increased oxygen supply by upscaling of the reaction vessel volume (Figure 6 B). The addition of glucose to the biotransformation buffer to improve cofactor regeneration was not beneficial. With these modified parameters a maximum concentration of around 1.1 mM (ca. 165 mg L^−1^ concentrated yeast cell suspension) *trans*-isopiperitenol could be obtained after 51 h of biotransformation, while a constant increase of *trans*-isopiperitenol could be observed over time.

**Figure 6:**
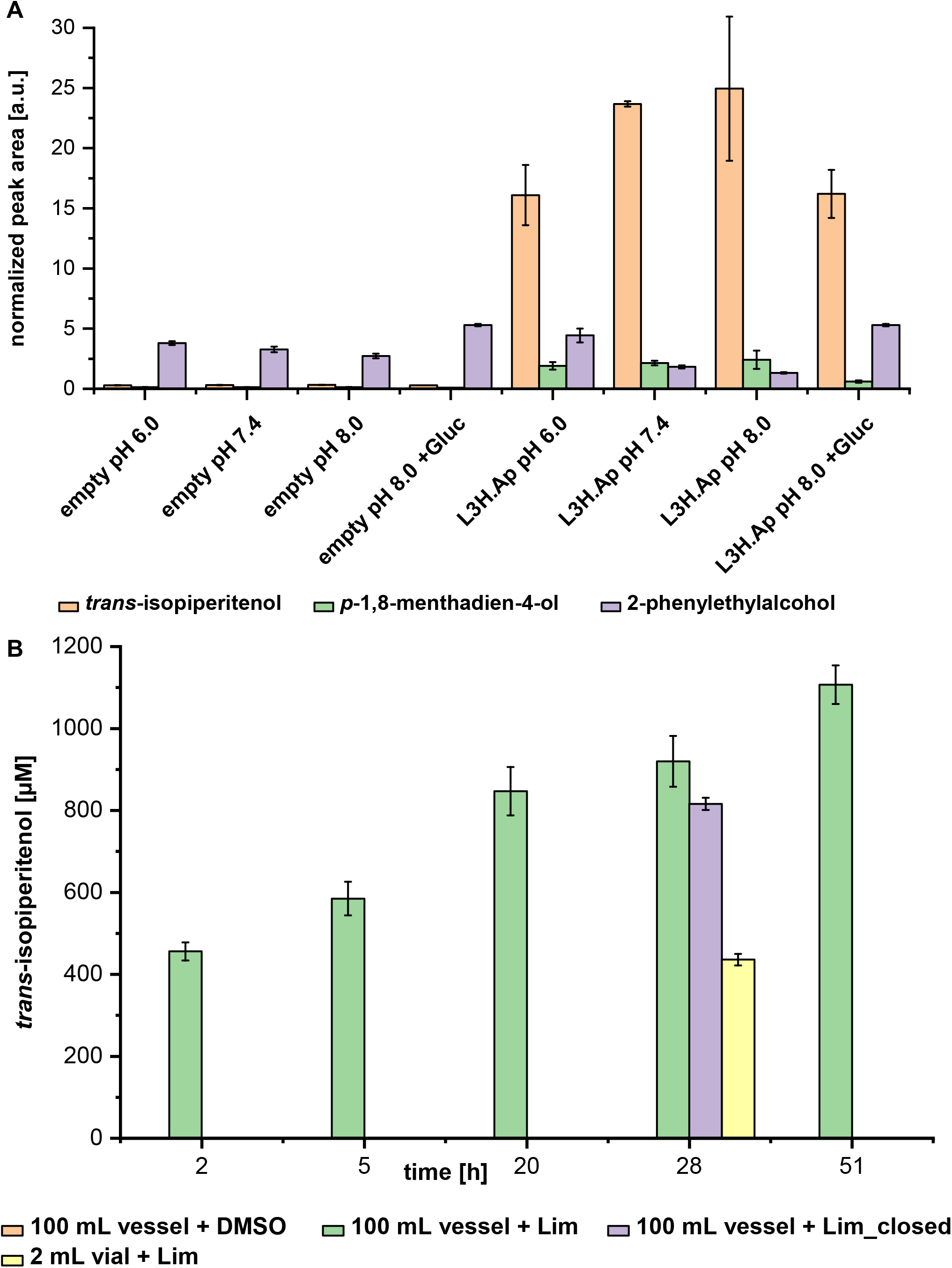
Product peak areas and *trans*-isopiperitenol concentrations achieved in (+)-limonene biotransformation experiments with *P. pastoris* X-33 strains expressing limonene-3-hydroxylase L3H.Ap under different conditions. Concentrated cell suspensions were used. Peak areas were determined by GC-MS analysis and normalized to the peak area of the internal standard (3-carene) and the number of cells corresponding to 225 OD units. *trans*-Isopiperitenol concentrations were calculated from normalized peak areas. The data points and error bars represent the mean values and standard deviations of three biological replicates (n = 3). (A) Investigation of the influence of buffer pH and glucose feeding on product formation. empty: *P. pastoris* strain with empty pBSYA1Z expression vector, L3H.Ap: strain expressing CYP65FA1 of *A. pullulans* from pBSYA1Z backbone. For biotransformation 100 mM potassium phosphate buffer with pH 6.0, 7.4 or 8.0 was used. For “+Gluc” the cell suspension contained 2 % glucose. When glucose was added, the pH of the biotransformation suspension decreased to under pH 5 during the experiment. (B) Investigation of the influence of reaction vessel volume on product formation at pH 8.0. For the experiment a *P. pastoris* strain expressing L3H.Ap from pBSYA1Z backbone was used. Biotransformation was carried out in reaction vessels with a total volume of 100 ml or 2 ml with cell suspension volumes of 7.5 mL or 750 μL, respectively. From “100 mL vessel + DMSO” and “100 mL vessel + Lim” samples were taken at different time points (2, 5, 20, 28 and 51 h after DMSO or limonene addition). The reaction vessels of “100 mL vessel + Lim_closed” and “2 mL vial + Lim” remained closed during the entire experiment and samples were only taken at 28 h.

### L3H.Ap can hydroxylate α-pinene, β-pinene and 3-carene

To further assess the substrate selectivity of L3H.Ap *in vivo*, other substrates were tested in biotransformation experiments (Table 1). While no products could be detected with γ-terpinene, α-phellandrene, toluene, 1-phenylethanol and 2-phenylethlyalcohol, we observed hydroxylation of 3-carene, α-pinene and β-pinene (Table 1, Figure 7, Figure S4 – Figure S7). The oxygenated monoterpene products were not, or only to a lesser extent, detected in experiments with the empty vector control.

**Table 1:** Overview of tested substrates and detected products of hydroxylation reactions with *P. pastoris* X-33 strain expressing L3H.Ap of *A. pullulans* CBS 100280 (EXF-150). For easier identification, the substances have been numbered in the respective figures (Fig. 2, 5, 7, S1 – S7). These numbers are given in brackets in the table behind the substance names.

| Substrate | Detected products after biotransformation with L3H.Ap in *P. pastoris* |
| --- | --- |
| (+)-limonene (1) | (+)-*trans*-isopiperitenol (2)<br>*p*-1,8-menthadien-4-ol (3) |
| (-)-limonene (4) | (-)-*trans*-isopiperitenol (5)<br>*p*-1,8-menthadien-4-ol (3)<br>carveol (6) |
| isopiperitenol ((+)-*trans*-/*cis*-mixture 2:1) | none |
| α-pinene (7) | verbenol (8)<br>verbenone (9)<br>further unknown products |
| β-pinene (10) | pinocarveol (11)<br>further unknown products |
| γ-terpinene | none |
| α-phellandrene | none |
| 3-carene (12) | *p*-1,5-menthadien-8-ol (13)<br>further unknown products |
| toluene | none |
| 1-phenylethanol | none |
| 2-phenylethylalcohol | none |
| DMSO | none |

**Figure 7:**
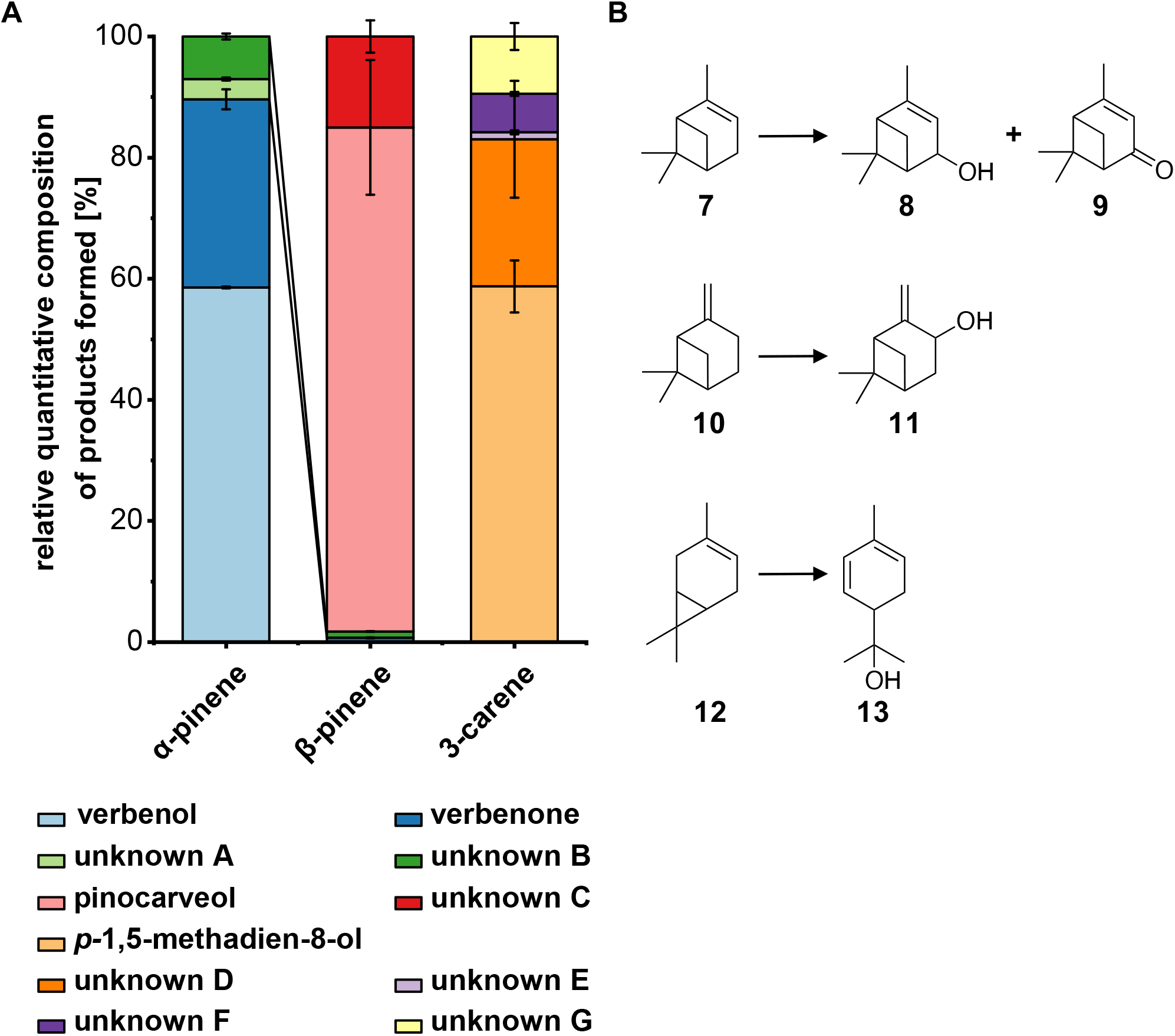
Hydroxylation of monoterpene substrates with *P. pastoris* X-33 expressing L3H.Ap. (A) Relative quantitative composition of products formed in biotransformation experiments of α-pinene, β-pinene or 3-carene with *P. pastoris* X-33 cells (concentrated cell suspension) expressing L3H.Ap from pBSYA1Z backbone. If possible, products were identified via comparison of retention times and mass-spectra with those of chemically synthesized reference substances, or by comparison of mass-spectra to those of the NIST mass spectral library (v14, MS spectra see Figure S4). Chromatograms see Figure S5 – Figure S7. The data points and error bars represent the mean values and standard deviations of three biological replicates (n = 3). (B) Substrates and main products of bioconversion reactions catalyzed by L3H.Ap. 7: α-pinene, 8: verbenol, 9: verbenone, 10: β-pinene, 11: pinocarveol, 12: 3-carene, 13: *p*-1,5-menthadien-8-ol.

## Discussion

In this work, we described two fungal CYP enzymes which can oxidize (+)-limonene at carbon atom 3, resulting in *trans*-isopiperitenol. Based on the protein sequence both enzymes, L3H.Ap and L3H.Hc, belong to the cytochrome P450 monooxygenase family of CYP65, which includes other fungal terpene hydroxylation enzymes described before (18).

In contrast to L3H.Ap and L3H.Hc, no limonene hydroxylation activity was detected for CYP65FA2.Ap, CYP65EX2.Ap and CYP65EW2.Ap as well as all other tested CYP candidate genes. This could either mean that these enzymes are unable to catalyze the desired reaction or that they are not functionally expressed in the yeast cells under the used conditions. In addition, CYP enzymes are known to require suitable reductase enzymes for their functionality, because these reductases provide the electrons necessary for the CYP reaction (19). Due to this dependency, it is also possible that no CYP activity could be measured because the CYP reductase provided in the yeast strains was not sufficient to enable the function of the tested CYP candidates. However, for overall CYP activity the redox partners can play a significant role (20). Furthermore, it is described that the type of co-expressed CYP reductase can even influence the product spectrum (21).

Although the physiological function of the limonene-3-hydroxylases in *A. pullulans*, *H. carpetanum* and other fungi is unclear, one can assume a role in modification/detoxification or utilization of monoterpenes produced by plants, which are colonized by such fungi (22). One would generally expect enzymes with high specific activity to carry out respective functions. The newly discovered limonene-3-hydroxylase from *A. pullulans* shows high stereospecificity and regioselectivity of the catalyzed reaction. Only small amounts (< 10 %) of the side product *p-*1,8-methadien-4-ol were produced from (+)-limonene by *P. pastoris* cells expressing L3H.Ap. Although some P450cam variants with selectivity preference for production of *trans*-isopiperitenol from (−)-limonene were described (6), the selectivity was lower than that of the fungal enzyme, if (+)-limonene was used as substrate (7). So far, the only highly specific limonene-3-hydroxylase enzymes are known to occur in *Mentha* plants (2, 10, 13). In peppermint, for example, PM17 catalyzes the key hydroxylation during the (−)-menthol biosynthesis pathway. In literature PM17 is described to convert (−)-limonene and (+)-limonene exclusively into the respective *trans*-isopiperitenols (23). In experiments with microsomal preparations from the epidermal oil glands (trichomes) of *Mentha* x *piperita* the (−)-isomer was favored by a factor of 2 over the (+)-enantiomer (24). Our results with *P. pastoris* cells indicate that the fungal L3H.Ap enzyme shows a reversed preference, which would be beneficial for the intended process using (+)-limonene from the citrus industry as starting material.

The titers of *trans*-isopiperitenol produced with *P. pastoris* cells are still low, reaching only up to around 165 mg L^−1^ of product in the concentrated cell suspension after around 51 h. Higher *trans*-isopiperitenol concentrations have already been reported for biotransformations with yeast cells expressing PM17 from peppermint (12). However, in our setup, using a different strain as well as a different CYP reductase, the *P. pastoris* strain expressing the fungal CYP enzyme L3H.Ap produced higher amounts of (+)-*trans-* isopiperitenol than the respective PM17-expressing strain. Since we did not investigate the protein concentrations in the cells, no conclusions about the differences of catalytic properties and substrate selectivity between the fungal enzymes and the peppermint limonene-3-hydroxylase are possible so far. In addition, as mentioned before, low CYP activity can also be caused by the absence of a suitable reductase enzyme. In order to be able to reliably compare the fungal enzymes with the plant protein, this interaction would also have to be optimized in further experiments, for example, by using a plant CPR in combination with PM17.

Apart from producing *trans*-isopiperitenol, it was found that L3H.Ap can also hydroxylate α- and β-pinene as well as 3-carene. The reactions delivered verbenol, verbenone, pinocarveol and *p*-1,5-menthadien-8-ol as main products. In the case of α-pinene, the oxidation of verbenol to verbenone might be either caused by L3H activity or by unspecific dehydrogenase activity present in yeast. The production of verbenol and verbenone from α-pinene in the same organism has already been described for bacteria as well as fungi and plant cells (6, 7, 25–28).

The activity of L3H.Ap on substrates other than limonene is in accordance with the conversion of α- and β-pinene by *Hormonema* sp. (4). However, while van Dyk and colleagues (1998) could also detect verbenol and verbenone as products of the α-pinene biotransformation experiment, β-pinene was converted to pinocamphone and 3-hydroxy-pinocamphone with *Hormonema* sp. (4). In addition to their experiments, oxidation of α- and β-pinene to several products, including verbenol, verbenone and pinocarveol, has been also reported for other fungi (4, 28, 29). Moreover, Chiu and colleagues (2019) recently described a CYP enzyme from mountain pine beetle that can hydroxylate α- and β-pinene as well as 3-carene (30). However, for this enzyme of the CYP6 family no activity for limonene isomers was detected. Furthermore, bacterial enzyme variants are known that can use carene, α-pinene or β-pinene as substrates, which are, however, converted into a variety of different products (7).

In this work we described the identification of two fungal limonene-3-hydroxylases that can hydroxylate (+)-limonene with high stereospecificity and regioselectivity into *trans*-isopiperitenol. In addition, further characterization of the L3H enzyme of *A. pullulans* showed that it can also convert (−)-limonene, 3-carene, α-pinene and β-pinene. By the use of yeasts expressing the CYP enzymes, novel catalysts for the conversion of (+)-limonene into (+)-*trans*-isopiperitenol could be provided that could become part of a (−)-menthol production process. Further investigations on their catalytic properties as well as optimization of cellular expression will be necessary to assess the full potential of the fungal CYP enzymes for biotechnological terpene conversions.

## Materials and methods

### Chemicals and media

All chemicals were purchased from Merck KGaA (Germany), previously Sigma-Aldrich (Germany) or Merck Millipore (Germany), or Carl-Roth (Germany) with different purities: (+)-limonene (≥ 99 %), (−)-limonene (≥ 99 %), α-pinene (≥ 98 %), (−)-β-pinene (≥ 99 %), γ-terpinene (≥ 98.5 %), α-phellandrene (≥ 95 %), 3-carene (≥ 98.5 %), 1-phenylethanol (≥ 98 %), 2-phenylethylalcohol (≥ 99 %), (−)-carveol (97 %), (−)-*trans*-pinocarveol (≥ 96 %), (*S*)-*cis*-verbenol (≥ 95 %), (1*S*)-(−)-verbenone (≥ 99 %). In addition, a 2:1 mixture of (+)-*trans*- and *cis*-isopiperitenol was obtained from the company Symrise (Germany).

Fungi, except *S. cerevisiae* and *P. pastoris,* were grown in yeast extract/malt extract (YM) medium (3 g L^−1^ yeast extract, 20 g L^−1^ malt extract, 10 g L^−1^ peptone, 10 g L^−1^ glucose) (31).

*S. cerevisiae* strains were cultivated in yeast extract peptone (YEP) medium (10 g L^−1^ yeast extract, 20 g L^−1^ peptone) or synthetic complete (SC) medium (6.7 g L^−1^ yeast nitrogen base without amino acids with ammonium sulfate, and amino acid supplements (32)). SC medium pH was adjusted to 6.3 with potassium hydroxide. If needed for selection of plasmids, medium without uracil and L-tryptophan was used. 20 g L^−1^ glucose were added as carbon source.

*P. pastoris* strains were cultured in YPD, YPDS, BMGY or BMMY media according to the *EasySelect™Pichia Expression Kit* manual (Invitrogen) and Weis *et al.* (2004) (33). *E. coli* was cultured in lysogeny broth (LB) medium (10 g L^−1^ tryptone, 5 g L^−1^ yeast extract and 5 g L^−1^ NaCl) or terrific broth (TB) medium (24 g L^−1^ yeast extract, 12 g L^−1^ tryptone, 5 g L^−1^ glycerol and 89 mM potassium phosphate buffer).

Antibiotics were added in the following concentrations, if required: ampicillin (Amp) 100 μg mL^−1^, Zeocin^TM^ (Zeo) 25 - 50 μg mL^−1^ (*E. coli)* or 100 – 500 μg mL^−1^ (*P. pastoris)*. For solid media 17 - 20 g L^−1^ agar-agar were added.

### Investigation of different fungi for limonene-3-hydroxylase activity

All strains tested for limonene-3-hydroxylase activity are listed in Table 2. In order to test the different Dothideomycetes for limonene-3-hydroxylase activities (Figure 2), 100 mL shake flasks with 25 ml YM medium were inoculated from 48 h old pre-culture and incubated for 48 h at 27 °C and 150 rpm (orbit: 2.5 cm). For the biotransformation experiment, 1 g L^−1^ (+)-limonene was added to the fungal cultures. 700 μL samples were taken from the reactions over a period of 5 hours (once per hour). Samples were extracted with one volume ethyl acetate and the *trans*-isopiperitenol amount was determined by GC-MS analysis. As negative control YM medium without cells was used.

**Table 2:**
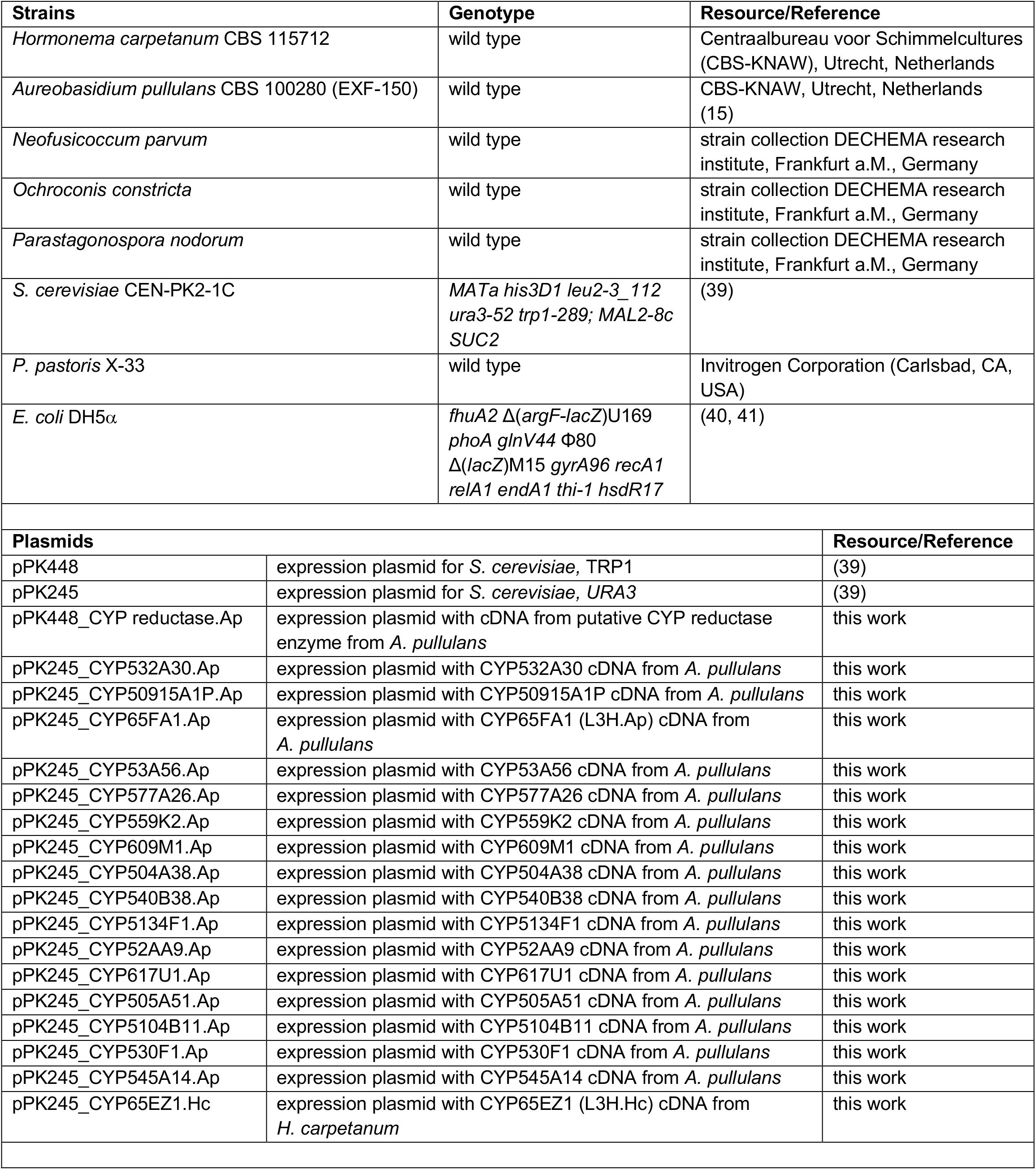
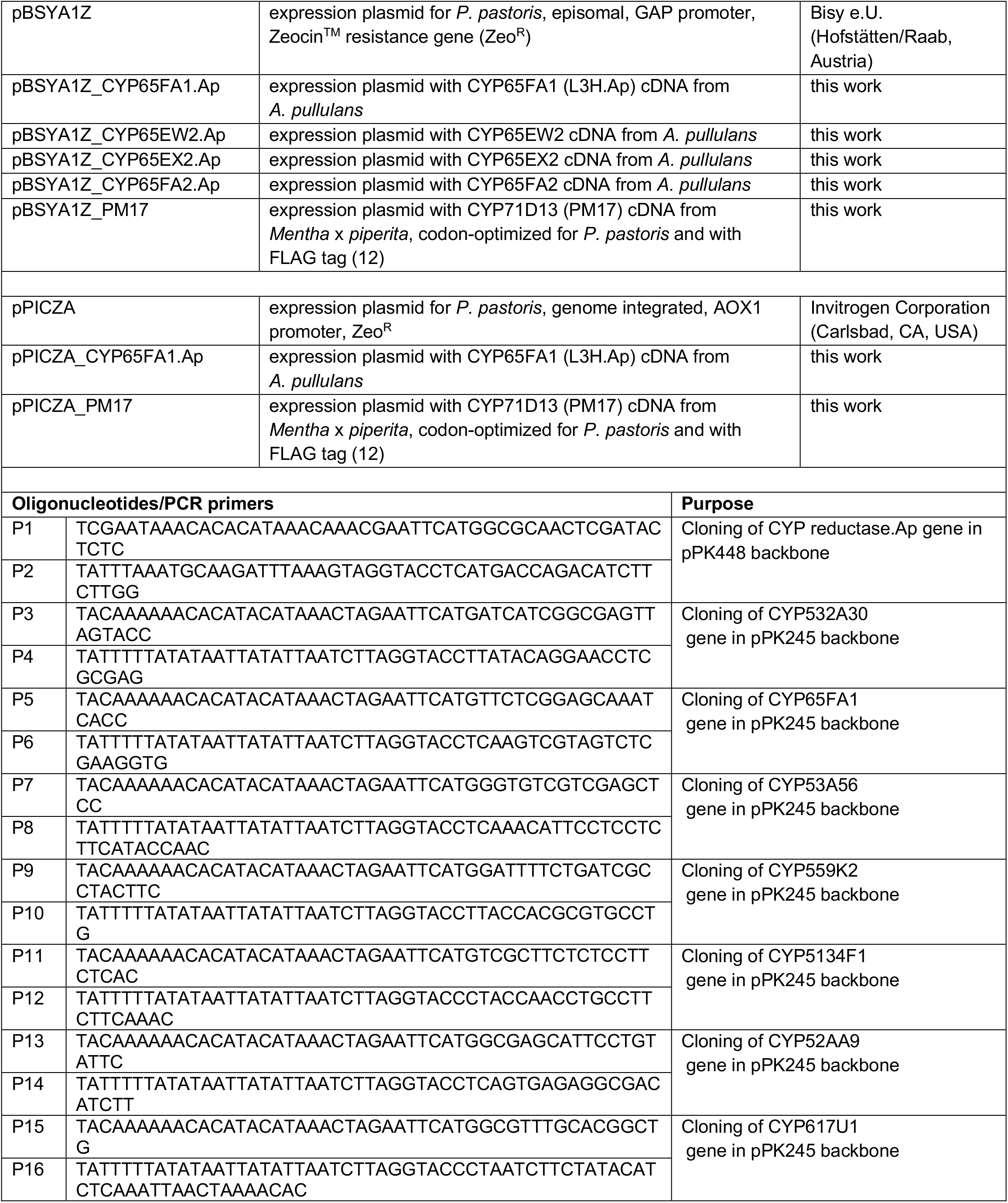
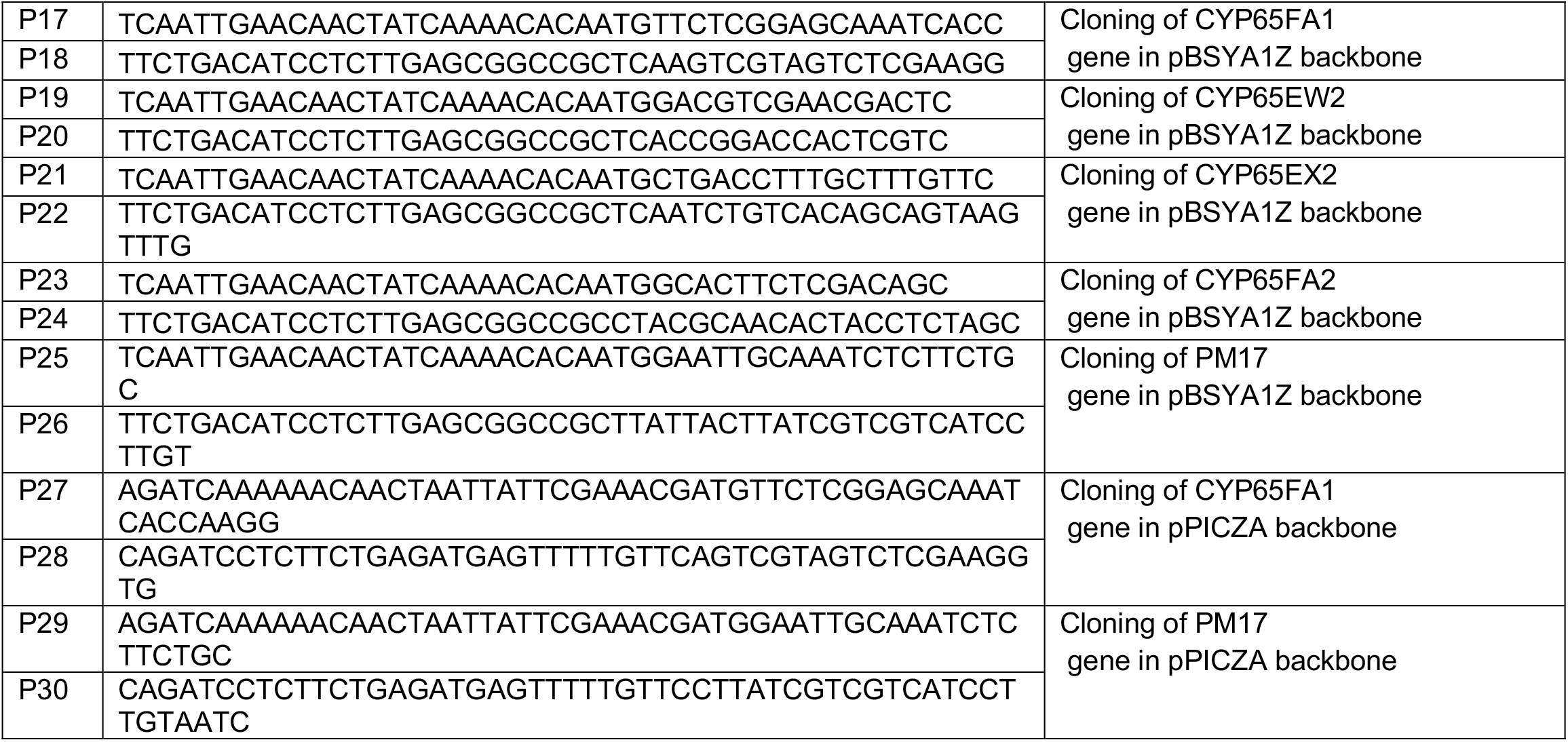
Fungal and bacterial strains, plasmids and oligonucleotides used in this study.

### Clustering of fungal cytochrome P450 monooxygenase sequences

The protein sequences of all 51 putative CYP enzymes from *A. pullulans* were clustered in a phylogenetic tree with the software *Geneious* (Biomatters Ltd., New Zealand) using default settings (multiple alignment: ClustalW alignment, cost matrix: BLOSUM, gap open cost: 10, gap extend cost: 0.1. Geneious tree builder: genetic distance model: Jukes-Cantor, tree build method: neighbor-joining, no outgroup, resampling method: bootstrap, number of replicates: 1,000).

### Construction of fungal CYP gene expression vectors and yeast strains

All yeast and *E. coli* strains, plasmids and oligonucleotides used in this study are listed in Table 2. cDNA sequences of selected CYP genes from *A. pullulans* EXF-150 (CBS 100280) were obtained through comparison between gene and protein sequence, stored in the NCBI database (accession numbers see Table S1). For synthesis of template cDNA, *A. pullulans* CBS 100280 cells were inoculated in 25 mL YM medium in a 100 mL shake flask. After incubation for 24 h at 28 °C and 130 rpm (orbit: 2.5 cm), 100 mg L^−1^(+)-limonene was added and incubated for further 24 h. Afterwards, cells were harvested, and total RNA isolated with *RNAeasy Plant MiniKit* (Qiagen) according to manufacturer's instructions. cDNA synthesis was performed with *iScript™ Select cDNA Synthesis Kit* (Biorad) using oligo(dT) primers. Afterwards, cDNAs of CYP genes were amplified via PCR. All expression plasmids used in this study were either constructed by the company BioCat GmbH (Germany) or constructed using Gibson isothermal assembly cloning (34, 35). *E. coli* cells were grown at 37 °C. Plasmids were isolated from *E. coli* using the *GeneJET Plasmid Miniprep Kit* (ThermoFisher Scientific). *S. cerevisiae* and *P. pastoris* strains were transformed with plasmids according to protocols of Gietz *et al.* (36, 37) or the *EasySelect™Pichia Expression Kit* manual (Invitrogen). Yeast cells were grown at 30 °C on SC selective medium or YPD containing Zeocin™.

### Investigation of fungal CYP candidates for limonene-3-hydroxylation in *S. cerevisiae*

For biotransformation experiments with *S. cerevisiae* a protocol based on publications by Emmerstorfer *et al.* was used (12, 38). *S. cerevisiae* CEN-PK2-1C strains, containing different pPK245 and pPK448 expression vectors (see Table 2), were grown on SC medium agar plates for 3 days. For initial testing of different CYP genes (Figure 4), main cultures were inoculated directly with a pin-head sized colony of yeast from agar plate in 50 mL SC medium in a 300 mL Erlenmeyer flask and incubated for a total of 72 h at 30 °C and 180 rpm (orbit: 2.5 cm). After 48 h, cells were fed with 1 % glucose. From stationary phase cultures 150 OD_600_ units were harvested (1,190 x g, 10 min) and resuspended in 500 μL of 100 mM potassium phosphate buffer (pH 7.4, total volume: 750 μl). Cell suspensions were then transferred into 2 mL glass vials and 300 mM (+)-limonene stock solution (in DMSO with 1 % Triton^®^ X-100) was added to yield a final limonene concentration of 6 mM. Glass vials were sealed loosely with screw caps and seal tape. Biotransformation was carried out for 24 h at 900 rpm (orbit: 2 – 3 mm) and 30 °C in a tabletop thermo shaker. Terpenoids were extracted with 250 μl methyl *tert*-butyl ether (MTBE) containing 100 μM 3-carene as internal standard. After phase separation by centrifugation for 10 min at 16,000 x g, organic layers were analyzed via GC-MS. CEN-PK2-1C strains containing the respective empty expression vectors were used as negative controls.

To quantify the produced *trans*-isopiperitenol amounts, stock solutions of isopiperitenol (2:1-mixture of *trans* and *cis*) with different concentrations were added to buffer aliquots of 750 μl and extracted in the same way with MTBE as the biotransformation samples.

### Biotransformation experiments with *Pichia pastoris*

An approach modified after Emmerstorfer *et al*. and Weis *et al.* was used (12, 33, 38). *P. pastoris* X-33 strains, containing different pBSYA1Z or pPICZA expression plasmids (see Table 2), were grown on YPD or BMDY medium agar plates containing 500 μg mL^−1^ Zeocin™ for 2 - 3 days at 30 °C. 25 mL BMDY medium in a 300 mL Erlenmeyer flask with 500 μg mL^−1^ Zeocin™ was inoculated directly with a pin-head sized colony of *P. pastoris* and incubated for around 24 h at30°C and 200 rpm (orbit: 2.5 cm). Afterwards 25 ml BMDY or BMMY medium was added to cultures with pBSYA1Z or pPICZA plasmids, respectively. 24 h later, 1 % glucose was added to strains with pBSYA1Z vectors. Strains with pPICZA plasmids were fed two or three times with 1 % methanol. 150 or 225 OD_600_ units from the 72 - 76 h old cultures were harvested (1,190 x g, 10 min) and resuspended in 375 or 500 μL of 100 mM potassium phosphate buffer (pH 6.0, 7.4 or 8.0). If glucose was added to the biotransformation reaction, 425 μL of buffer and 75 μL of a 20 % glucose solution were used. Cell suspensions were then transferred into 2 mL glass vials and 300 mM (+)-or (−)-limonene stock solution (in DMSO with 1 % of Triton^®^ X-100) was added to yield a final concentration of 6 mM limonene. Glass vials were sealed loosely with screw caps and seal tape. Biotransformation was carried out for 24 - 28 h at 900 rpm (orbit: 2 – 3 mm) at 30°C in a tabletop thermo shaker. Terpenoids were extracted with 250 μL MTBE containing 100 μM 3-carene as internal standard. After phase separation by centrifugation for 10 min at 16,000 x g, organic layers were analyzed via GC-MS. For experiments in 100 mL reaction vessels, for each biotransformation batch 2250 OD_600_ units were harvested (1,190 x g, 10 min) and resuspended in 5 mL of 100 mM potassium phosphate buffer (pH 8.0). The cell suspensions were transferred to 100 mL glass bottles and limonene stock solution was added to a final concentration of 6 mM limonene. The vessels were tightly closed with screw caps and Parafilm^®^. The glass bottles were incubated for 51 h in a cabinet incubator at 30 °C and 250 rpm (orbit: 2.5 cm). Samples of 750 μl were taken at 2, 5, 20, 28 and 51 h and extracted as described for the other *Pichia* samples. Some batches were kept closed throughout the experiment without sampling to make sure that no *trans*-isopiperitenol evaporates. *P. pastoris* strains containing the respective empty expression vectors were used as negative controls in all experiments.

To quantify the produced *trans*-isopiperitenol amounts, stock solutions of isopiperitenol (2:1-mixture of *trans* and *cis*) with different concentrations were added to cell suspension aliquots of a *P. pastoris* strain, containing the empty pBSYA1Z vector, and extracted in the same way with MTBE as the biotransformation samples.

For the bioconversion experiments with substrates other than (+)- and (−)-limonene, the different substances (see Table 1) were dissolved in DMSO with 1 % of Triton^®^ X-100 and added to the biotransformation reactions to give a final concentration of 6 mM. Only in the case of isopiperitenol (2:1-mixture of *trans* and *cis*) 0.6 μM were used. Biotransformation samples were extracted as described, but without internal standard.

### GC-MS analysis

For GC-MS analysis of fungi biotransformation samples, except *S. cerevisiae* and *P. pastoris,* a GC-17A QP5050A system (Shimadzu) with a VB-5 column (30 m x 0.25 mm x 0.25 μm, ValcoBond^®^, Valco Instruments Co. Inc. and VICI AG) or a GCMS-QP2010 SE system (Shimadzu) containing a DB-5 column (30 m x 0.25 mm x 0.25 μm, Agilent) was used. Measurements were conducted as follows: helium as carrier gas, split ratio 10, injection at 250 °C with a sample volume of 1 μl, a column flow of 1.2 mL min^−1^ and a linear velocity of 39.2 cm s^−1^. The column temperature was programmed as follows: 60 °C for 3 min, with 10 °C min^−1^ up to 240 °C followed by 3 min at 240 °C.

Samples of biotransformation experiments with *S. cerevisiae* and *P. pastoris* were analyzed by a GCMS-QP2010 SE (Shimadzu) system containing a DB-5 column (30 m x 0.25 mm x 0.25 μm, Agilent). Measurements were conducted as follows: helium as carrier gas, split ratio 1 or 5, injections at 270 °C, column flow of 0.7 mL min^−1^ and a linear velocity of 30 cm s^−1^. The column temperature was programmed as follows: 40 °C, 9 °C min^−1^ up to 175 °C followed by 30 °C min^−1^ up to 300 °C.

*trans*-Isopiperitenol, carveol, verbenol, verbenone and pinocarveol were identified by comparison of retention times and mass-spectra with those of chemically synthesized reference substances (see Chemicals and media). *p*-1,8-menthadien-4-ol and *p*-1,5-menthadien-8-ol and other products were identified by comparison of mass-spectra to those of the NIST mass spectral library (v14) or the *MassLib* mass spectral library (Symrise Holzminden, V9.4 0). Identity was assumed if the similarity index was equal to or higher than 90 %. Absolute concentrations of *trans*-isopiperitenol were calculated from chromatogram peak areas by comparison to a calibration curve prepared by measuring a dilution series of isopiperitenol reference substance with known concentrations.

## Acknowledgments

We thank Prof. David R. Nelson for assigning unique identifiers to the CYP proteins. This research was supported by funds of the Federal Ministry of Food and Agriculture (BMEL) based on a decision of the Parliament of the Federal Republic of Germany via the Fachagentur Nachwachsende Rohstoffe e.V. (FNR, FKZ 22030615) under ERA-NET, funded from the European Union’s Seventh Research Framework Program (ERA IB-2, 6th call, project BioProMo), by funds of the Federal Ministry of Education and Research (BMBF, FKZ 0315810, BioIndustrie2021) and by the LOEWE project AROMAplus of the State of Hessen (Germany).

## Author Contributions

F.M.S, I.S., M.M.W.E., H.S., J.S. and M.B designed the research. F.M.S, I.S., M.M.W.E. and E.B. performed the experiments. J.P. contributed reagents and analysis tools. F.M.S, I.S., M.M.W.E., J.P. and M.B. analyzed the data. F.M.S. and M.B. wrote the manuscript with input from all authors.

We declare we have no conflicts of interest regarding the content of this article.

